# Heterologous expression of methylxanthine synthesis enzymes in mammalian cells and their use as reporter proteins

**DOI:** 10.1101/2020.12.29.424659

**Authors:** Brandon T. Cisneros, Neal K. Devaraj

## Abstract

This work demonstrates the reconstitution of active methylxanthine synthesis enzymes in human cells and their potential use as inducible reporter enzymes. A variety of plant enzymes involved in caffeine synthesis have been characterized *in vitro* and several of these methylxanthine synthesis enzymes have been heterologously-expressed in yeast or bacteria. In this work, enzymes from *Coffea arabica, Camellia sinensis*, and *Paullinia cupana* have been heterologously-expressed in human cells. We demonstrate that the enzymes tested exhibit similar patterns of activity with a set of xanthine substrates in human cells when compared to previous reports of *in vitro* activity. We demonstrate that the activity of these enzymes can be used as a reporter for juxtacrine signaling using synNotch-induced expression in the presence of an appropriate substrate. When used in combination with synthetic caffeine receptors, this work has potential for use as an *in vivo* reporter (e.g. enabling non-invasive monitoring of cell-cell interactions after a cellular transplant) or in synthetic intercellular signaling a methylxanthine, such as caffeine, acting as a synthetic paracrine hormone.

## Introduction

Xanthine is ubiquitous in life as a degradation product of the purine bases found in DNA. A variety of methylxanthines can be produced by N-methylation at positions 1, 3, and 7 to form seven possible methylxanthine products: three monomethylxanthines, three dimethylxanthines, and 1,3,7-trimethylxanthine (caffeine, 137mX) (**Figure 1**). Several methylxanthines have notable pharmacologic properties that are of relevance to humans: theophylline, theobromine, and caffeine. Theophylline and its derivatives are used in medicine for their effects on the respiratory system or as stimulants.[1] Theobromine is a mild stimulant found in chocolate.[2] Caffeine is a stimulant drug and is by far the xanthine derivative of most interest to people. It is consumed by roughly 85% of Americans on a daily basis[3] and the cultivation of caffeine-containing plants is of immense economic value in the present day and historically.[4]

**Figure 1:**
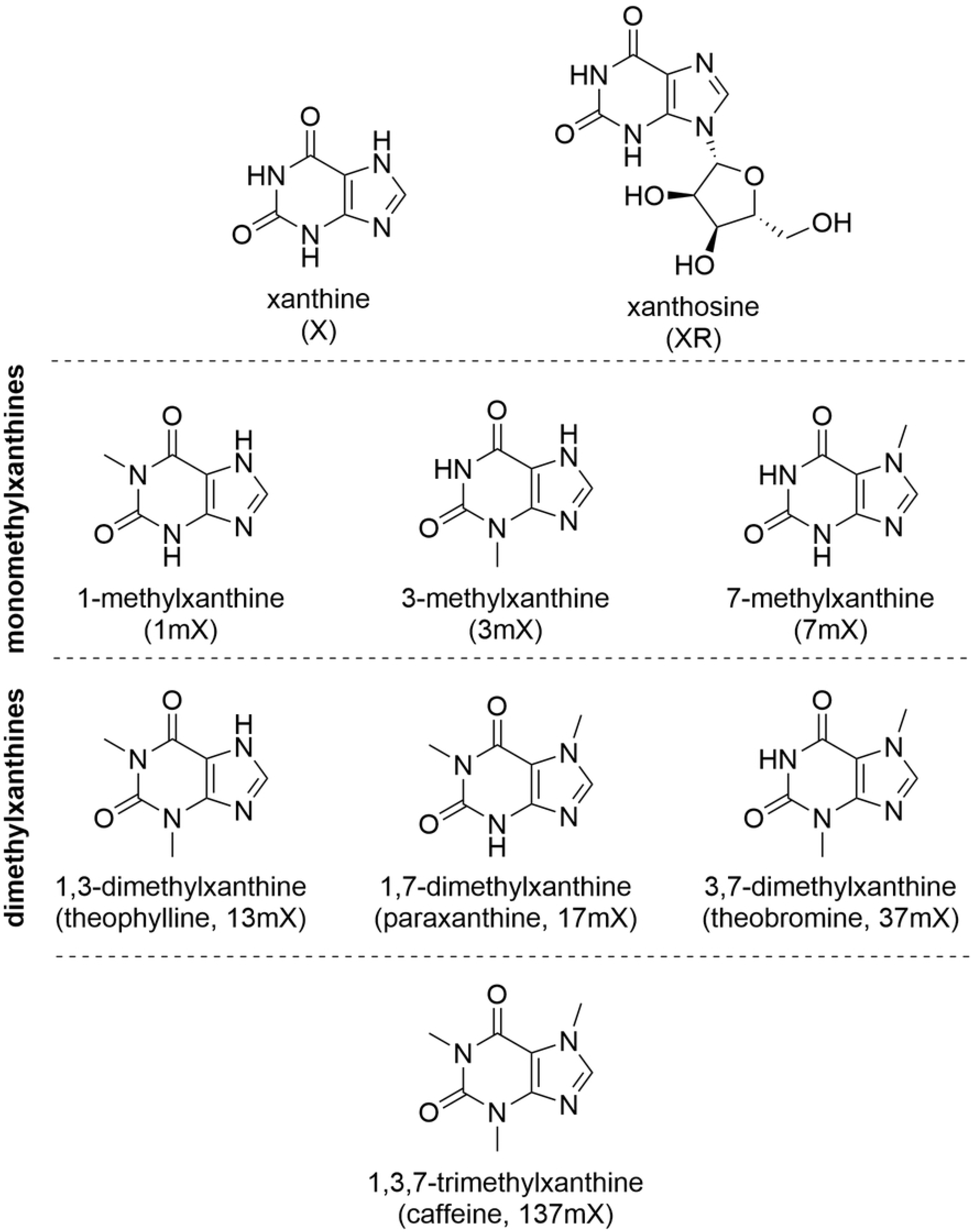
Xanthine derivatives used in caffeine biosynthesis pathways. Structures of the xanthine, xanthosine, and all possible methylxanthine substitution patterns.

A majority of caffeine consumed by people is derived from coffee (*Coffea spp.*, namely *Coffea arabica*) and tea (*Camellia sinensis*) with a small contribution from other plants like guarana (*Paullinia cupana*) and yerba mate (*Ilex paraguariensis)*.[5] The dominant enzymes and corresponding genes involved in the biosynthesis of caffeine have been identified and characterized for coffee, tea, and guarana.[6–13] More recently, genome sequencing data has become available and is being used to build a more thorough understanding of the multitude of enzymes involved in caffeine biosynthesis.[13–19]

In both coffee and tea, caffeine synthesis pathways (**Figure 2A**) can convert xanthosine to caffeine via three sequential steps: N-7 methylation (with loss of the ribose moiety), N-3 methylation, and N-1 methylation. In coffee, this process is carried out by CaXMT1 (*Coffea arabica* xanthosine methyltransferase 1, BAB39215), CaMXMT (*Coffea arabica* methylxanthine methyltransferase 1, XP_027086104), and CaDXMT (*Coffea arabica* dimethylxanthine methyltransferase, BAC75663). Coffee additionally contains the multi-functional enzyme CCS1 (coffee caffeine synthase 1, BAC43760.1) which can perform the latter two methylations, but this enzyme does not appear to be primarily responsible for caffeine production in coffee.[11] In tea, TCS2 (*Camellia sinensis* caffeine synthase 2, AB031281) carries out the N-7 methylation and TCS1 (*Camellia sinensis* caffeine synthase 1, AB031280) carries out both of the successive N-3 and N-1 methylations.[13] These methylating enzymes all stoichiometrically consume S-adenosylmethionine (SAM) as a methyl donor and produce S-adenosylhomocysteine (SAH).

**Figure 2:**
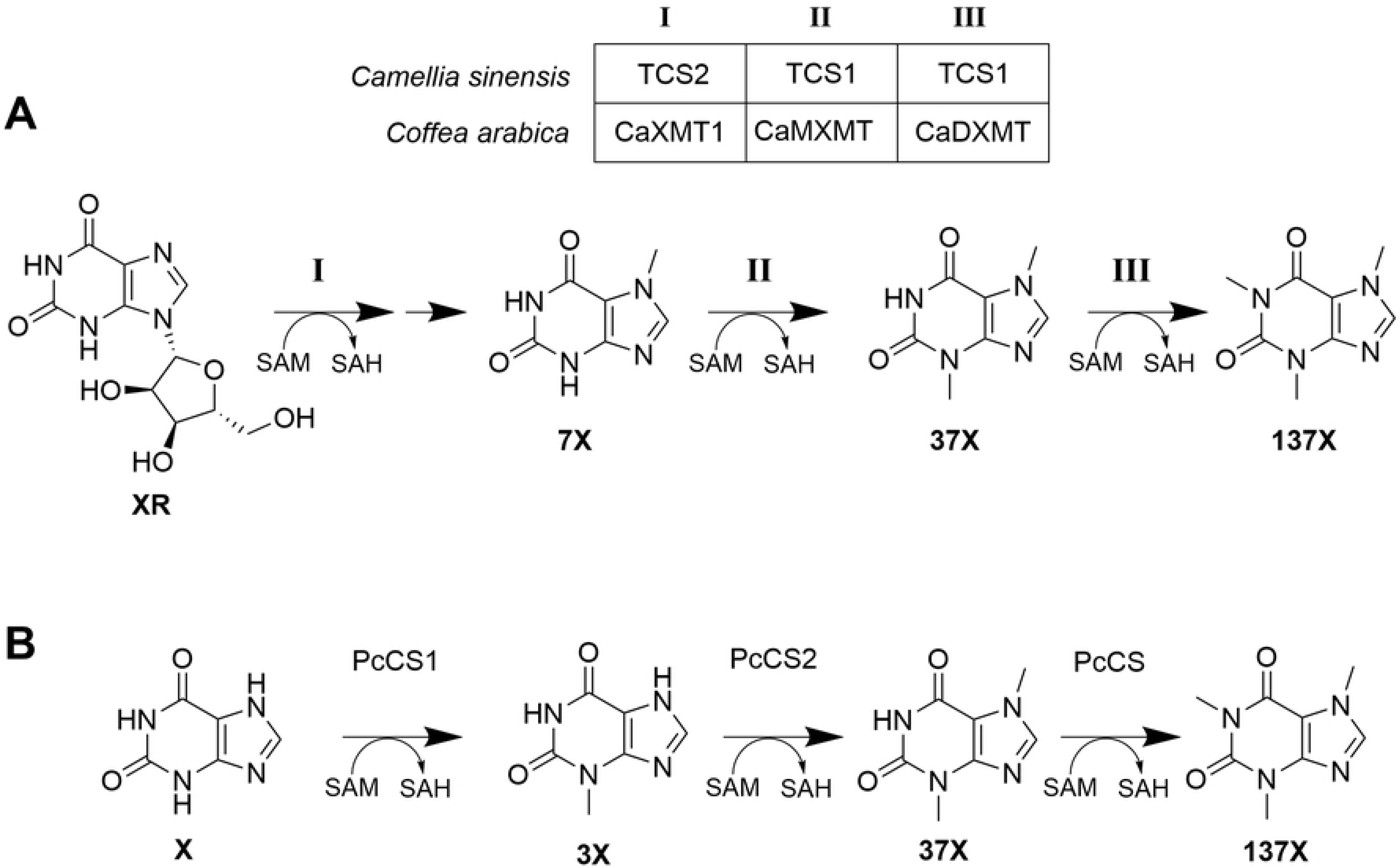
Caffeine biosynthesis pathways. (A) Pathway for biosynthesis of caffeine in tea and coffee. The cleavage of ribose from 7-methylxanthosine after step I is not shown, as the mechanistic details are yet unknown. (B) Pathway for the biosynthesis of caffeine in guarana

The caffeine synthesis pathway in guarana produces caffeine in a different manner than coffee or tea (**Figure 2B**). In guarana, xanthine is used as a substrate instead of xanthosine, and the three methylation steps take place in a different order: N-3, N-7, followed by N-1 methylation. These methylation steps are carried out by *Paullinia cupana* caffeine synthase 1 (PcCS1, EC766748), *Paullinia cupana* caffeine synthase 2 (PcCS2, EC778019), and *Paullinia cupana* caffeine synthase (PcCS, BK008796), respectively.[13] As with coffee and tea, these enzymes consume SAM as a methyl donor and produce SAH.

Previous work has demonstrated that a hybrid coffee-tea caffeine synthesis pathway using CaXMT1 and TCS1 can be used to produce caffeine de novo in *E. coli* and *S. cerevisiae.* Subsequent work has expanded on this work to optimizing production of a variety of other methylxanthines in *S. cerevisiae.* Another effort focused on increasing metabolic flux through purine degradation pathways to increase the yields of de novo caffeine synthesis. The fact that multiple caffeine synthesis pathways had been successfully reconstituted in *E. coli* and *S. cerevisiae* combined with the relatively high activity of these enzymes suggested that it may be feasible to reconstitute them in mammalian cells as well.

If these enzymes were active in mammalian cells, we supposed that they may be a useful candidate for use in orthogonal signaling schemes. Four important criteria[20] for viable approaches have been previously identified: (i) the signal molecule should be freely diffusible; (ii) detection should be highly specific; (iii) the signal molecule should be produced from endogenous components; (iv) the number of enzymatic transformations needed to make it should be minimal. Criteria i, iii, and iv seem to be clearly met by caffeine and the other methylxanthines. Criterion ii is satisfied by the existence of synthetic caffeine receptors.[21] The combination of methylxanthine synthesis enzymes with synthetic caffeine receptors would likely be suitable for orthogonal intercellular signaling.

We set out to determine if methylxanthine synthesis enzymes from caffeine synthesis pathways in plants are functional when heterologously-expressed in mammalian cells. We found that, except for PcCS, all the methylxanthine synthesis enzymes tested showed the same patterns of activity in mammalian cells as they show *in vitro*. We sought to test whether a methylxanthine synthesis enzyme could be used as a reporter for gene expression. We demonstrated that production of caffeine from paraxathine by CCS1 could be used as a reporter for juxtacrine cell signaling. We believe that this work has relevance to the field of engineered intercellular signaling and has a potential application in enabling the use of caffeine as a small molecule reporter for monitoring cellular activity *in vivo*.

## Results and discussion

### Methylxanthine synthesis enzymes are active in mammalian cells

Several methylxanthine synthesis enzymes from three plants were chosen to potentially express in mammalian cells: CaXMT1, CaMXMT, CaDXMT, CCS1 from *Coffea arabica*; TCS1 and TCS2 from *Camellia sinensis*; and PcCS from *Paullinia cupana.* The dominant reactions performed by these enzymes are shown in **Table 1**. Two putative ancestral enzymes CamelliaAncCS (CsAncCS, MW309842) and PaulliniaAncCS2 (PcAncCS2, MW309843) were chosen as well. These latter two enzymes were chosen because previous work suggested that they may possess the ability to produce some quantity of paraxanthine from 7-methylxanthine,[22] a property not exhibited by any known natural methylxanthine synthesis enzymes.

**Table 1:**
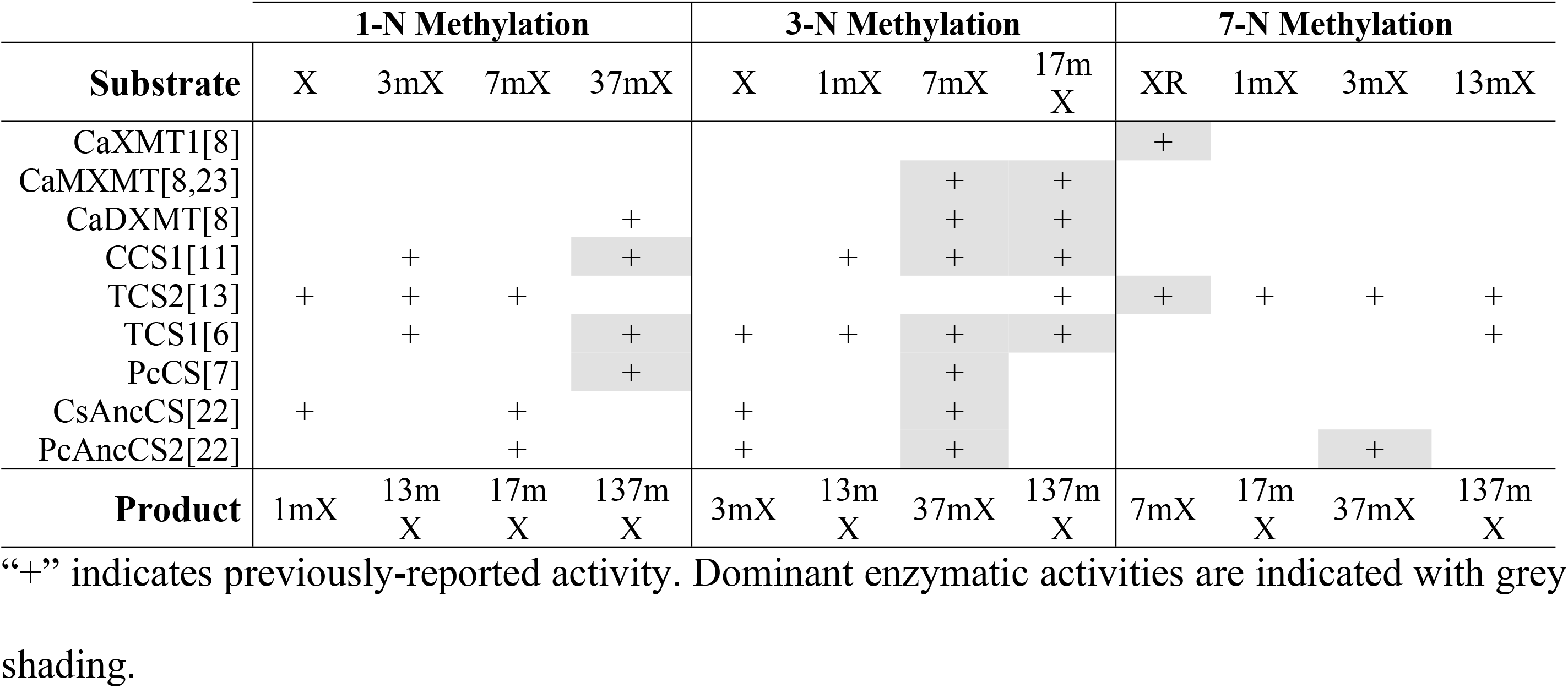
Expected Methylation of Xanthine Substrates by Methyltransferase Enzymes.

| Substrate | 1-N Methylation |  |  |  | 3-N Methylation |  |  |  | 7-N Methylation |  |  |  |
| --- | --- | --- | --- | --- | --- | --- | --- | --- | --- | --- | --- | --- |
|  | X | 3mX | 7mX | 37mX | X | 1mX | 7mX | 17mX | XR | 1mX | 3mX | 13mX |
| CaXMT1[8] |  |  |  |  |  |  |  |  | + |  |  |  |
| CaMXMT[8,23] |  |  |  |  |  |  | + | + |  |  |  |  |
| CaDXMT[8] |  |  |  | + |  |  | + | + |  |  |  |  |
| CCS1[11] |  | + |  | + |  | + | + | + |  |  |  |  |
| TCS2[13] | + | + | + |  |  |  |  | + | + | + | + | + |
| TCS1[6] |  | + |  | + | + | + | + | + |  |  |  | + |
| PcCS[7] |  |  |  | + |  |  | + |  |  |  |  |  |
| CsAncCS[22] | + |  | + |  | + |  | + |  |  |  |  |  |
| PcAncCS2[22] |  |  | + |  | + |  | + |  |  |  | + |  |
| **Product** | 1mX | 13mX | 17mX | 137mX | 3mX | 13mX | 37mX | 137mX | 7mX | 17mX | 37mX | 137mX |

“+” indicates previously-reported activity. Dominant enzymatic activities are indicated with grey shading.

We constructed expression vectors for each of the methylxanthine synthesis enzymes using the Sleeping Beauty transposon vectors pSBBi-RP and pSBBI-GB.[24] HEK-293T human embryonic kidney cells were transduced using the Sleeping Beauty transposase system by co-transfection with pCMV(CAT) T7-SB100[25–27] and selected with an appropriate antibiotic to generate a stable cell line expressing each of the chosen enzymes. To test the enzyme activity in cells, we first seeded each cell line in a 12-well plate. After 24 h, they were then exposed to a fixed concentration (200 μM) of each of the following substrates: xanthine (X), 1-methylxanthine (1mX), 3-methylxanthine (3mX), 7-methylxanthine (7mX), 1,3-dimethylxanthine (13mX, theophylline), 1,7-dimethylxanthine (17mX, paraxanthine), 3,7-dimethylxanthine (37mX, theobromine), and xanthosine (XR) (for enzymes with expected activity against this substrate). This concentration of substrate was chosen to be sufficiently high that the dominant enzyme activity for each enzyme would be likely to provide a detectable yield of product at the end of the experiment without being so high as to significantly slow cell growth or cause toxicity. After 72 h of growth, the supernatant was harvested and analyzed by high performance liquid chromatography (HPLC) with a UV-Vis diode array detector. Product identity was confirmed by comparison to genuine standards. No product formation was observed for any substrate with untransformed HEK-293T cells. A representative example of the HPLC chromatograms is shown in **Figure 3** for TCS1-expressing cells. Other HPLC chromatograms are available in **S1 File**.

**Figure 3:**
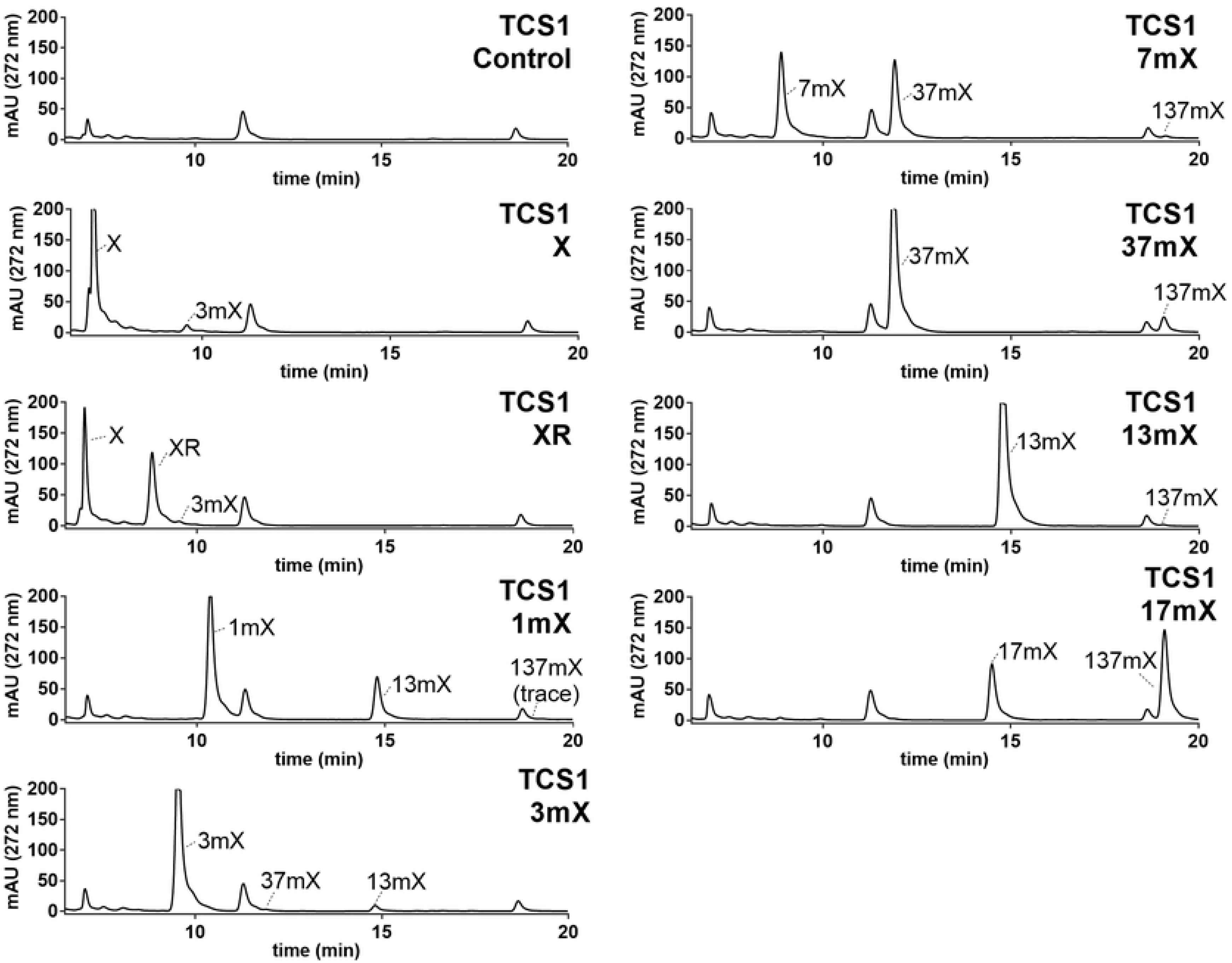
Representative chromatograms for cell culture with xanthine substrates. HPLC chromatograms of TCS1-expressing HEK-293T cells grown in 200 μM of each of the xanthine substrates for 72 h with relevant xanthine peaks identified

We found that the observed activities (**Table 2**) of the methylxanthine synthesis enzymes tested to be largely consistent with previous reports **(Table 1)** of their activities *in vitro*. Although these experiments were not designed to provide a quantitative measurement of enzyme activity, comparison of the relative activities between substrates for a given enzyme also reflect previously-reported enzyme activities. The largest discrepancy between our observations and expectations was with the lack of any observed activity for PcCS-expressing cells. Under the reaction conditions, we expected to detect at least trace product formation when 37mX or 7mX were used as substrates, but no product was observed in either case. It is possible that there were enzyme-specific problems that inhibited its expression or activity under the experimental conditions, but this was not investigated further. All other discrepancies we noted were for cases where small quantities of product were expected (i.e. for the off-target reactions of TCS2) or where the enzymes had not been thoroughly studied (i.e. for CsAncCS and PcAncCS2). These findings are likely due to differences in the sensitivity of the methods we used compared to those used in previous studies.

**Table 2:**
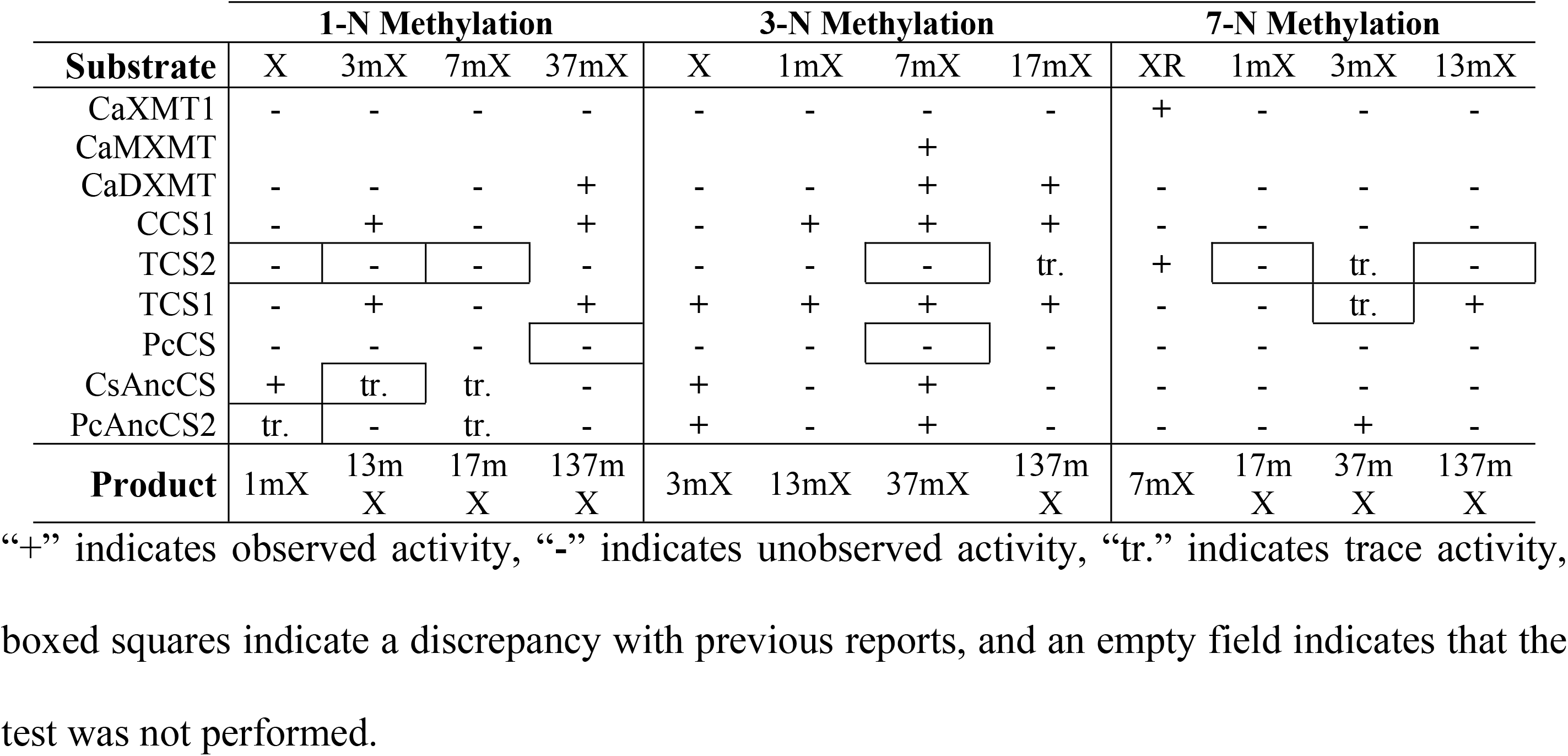
Observed Methylation of Xanthine Substrates by Methyltransferase Enzymes.

| Substrate | 1-N Methylation |  |  |  | 3-N Methylation |  |  |  | 7-N Methylation |  |  |  |
| --- | --- | --- | --- | --- | --- | --- | --- | --- | --- | --- | --- | --- |
|  | X | 3mX | 7mX | 37mX | X | 1mX | 7mX | 17mX | XR | 1mX | 3mX | 13mX |
| CaXMT1 | - | - | - | - | - | - | - | - | + | - | - | - |
| CaMXMT |  |  |  |  |  |  | + |  |  |  |  |  |
| CaDXMT | - | - | - | + | - | - | + | + | - | - | - | - |
| CCS1 | - | + | - | + | - | + | + | + | - | - | - | - |
| TCS2 | - | - | - | - | - | - | - | tr. | + | - | tr. | - |
| TCS1 | - | + | - | + | + | + | + | + | - | - | tr. | + |
| PcCS | - | - | - | - | - | - | - | - | - | - | - | - |
| CsAncCS | + | tr. | tr. | - | + | - | + | - | - | - | - | - |
| PcAncCS2 | tr. | - | tr. | - | + | - | + | - | - | - | + | - |
| Product | 1mX | 13mX | 17mX | 137mX | 3mX | 13mX | 37mX | 137mX | 7mX | 17mX | 37mX | 137mX |

“+” indicates observed activity, “-” indicates unobserved activity, “tr.” indicates trace activity, boxed squares indicate a discrepancy with previous reports, and an empty field indicates that the test was not performed.

### Xanthosine is degraded to xanthine in mammalian cells

While chromatograms were readily interpretable for most substrates, the chromatograms for XR were more complicated to interpret for several reasons. For substrates other than XR, the largest peak observed on the chromatogram is from the substrate itself or the product made by methylation of that substrate. As shown in **Figure 3**, when XR was supplied as a substrate to TCS1-expressing cells, a small quantity of 3mX was observed even though TCS1 is not expected to methylate XR. In addition, the most prominent peak observed is for xanthine rather than xanthosine. This can be explained by xanthosine being degraded to xanthine as part of normal purine degradation pathways.[28] If xanthine is being formed in situ, then we would expect to see 3mX being synthesized via the known action of TCS1 on X. Indeed, we do observe evidence of this.

### Production of 7mX from XR by TCS2- and CaXMT1-expressing cells can be verified by co-culture with CCS1-expressing cells

Because 7mX and XR have nearly-identical retention times (as seen in Figure 3), we could not use chromatography alone to definitively determine if TCS2 and CaXMT1 produce 7mX from XR in mammalian cells. Our experiments with CCS1-expressing cells showed that they rapidly produced 37mX when supplied with 7mX and do not produce any products when supplied with X or XR. We hypothesized that if we co-cultured TCS2 or CaXMT1 cells with CCS1 cells in the presence of XR then some portion of the 7mX produced would be converted to the easily-separable 37mX. This would allow us to know with certitude that 7mX was indeed being produced by TCS2 or CaXMT1 from XR in situ. To do this, we compared two populations of cells grown for 72h in 400 μM XR: TCS2- or CaXMT1-expressing cells alone and a one-to-one co-culture of CCS1-expressing cells and TCS2- or CaXMT1-expressing cells.

As shown in **Figure 4**, TCS2-expressing cells grown alone produce a broad, complex peak composed of XR and 7mX. When TCS2-expressing cells are co-cultured with CCS1-expressing cells, the intensity of that complex peak decreases and a 37mX peak appears. The same scenario occurred for CaXMT1. These experiments demonstrate that the inseparable peak occurring in the chromatograms for TCS2 and CaXMT1 definitively contains 7mX, as the only way CCS1 can produce 37mX is by methylation of 7mX. In addition, these results demonstrate the possibility of reconstituting a multi-step methylxanthine synthesis pathway in mammalian cells by co-culture of cells in which separate cells each perform a single step of the process.

**Figure 4:**
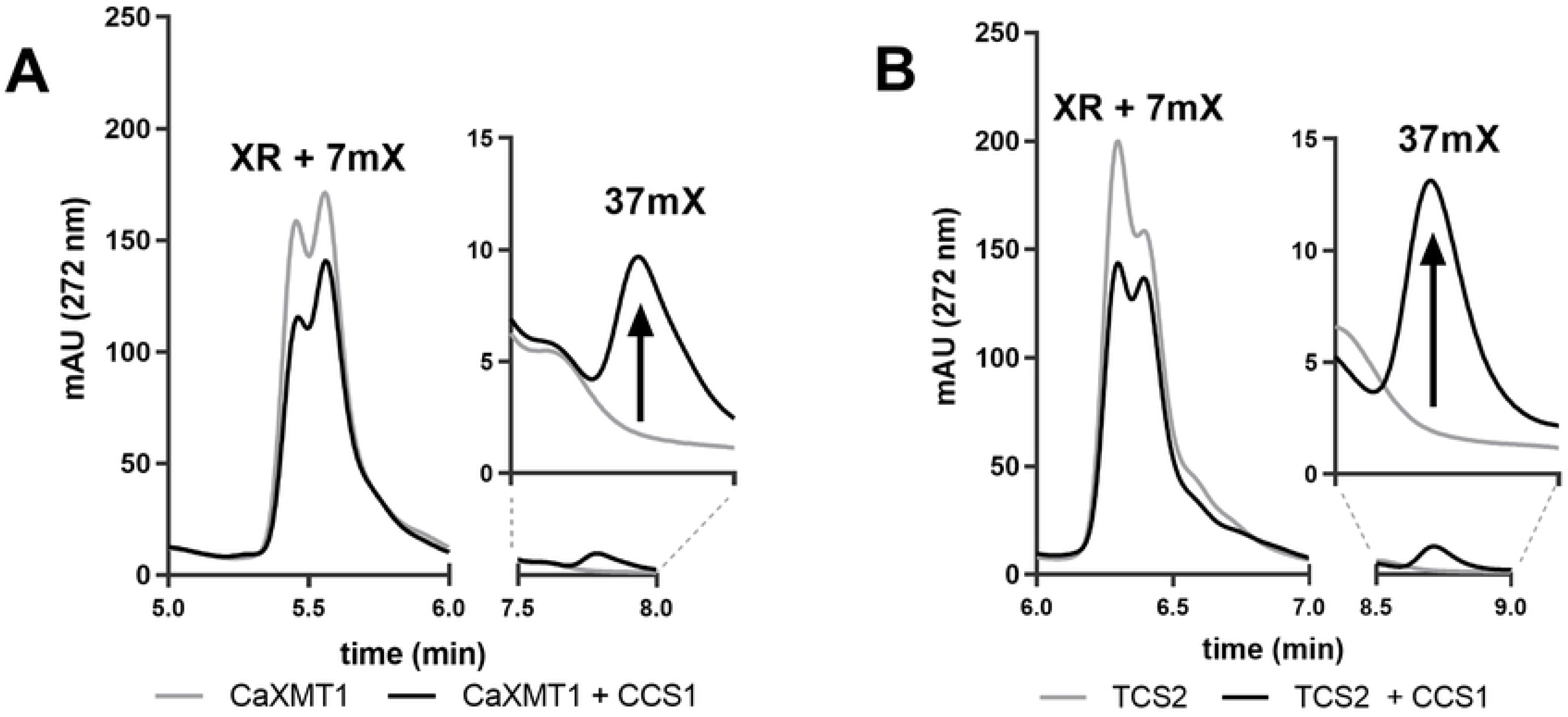
HPLC chromatograms demonstrating definitive production of 7mX by CaXMT1 and TCS2. 37mX is produced after co-culture with CCS1-expressing cells indicating definitive production of 7mX from XR by either enzyme. This is shown in (A) CaXMT-1 expressing cells and (B) TCS2-expressing cells. Inset in each shows detail of the 37mX peak

### CsAncCS produces 1mX and 3mX from endogenous xanthine

In our data collected with CsAncCS, we observed small quantities of 1mX and 3mX throughout all samples tested, including the negative control (**Figure 5**). We first assumed this could have been the result of contamination, but this observation persisted when the experiment was repeated. Although present only in trace amounts in all the samples, the relative size of these peaks appeared to diminish when 7mX was provided to the cells as a substrate. Previous work has reported that CsAncCS has a K_M_ = 26.7 μM with 7mX.[22] As such, 200 μM of 7mX could be competitively inhibiting the activity of CsAncCS upon endogenous X and limiting the production of 1mX and 3mX. Observing detectable product formation from endogenous xanthine alone suggests that CsAncCS may be a useful enzyme as part of a pathway for de novo methylxanthine synthesis in cells, mammalian or otherwise. This could be further enhanced by increasing molecular flux through the purine degradation pathways to increase endogenous xanthine levels.[29,30]

**Figure 5:**
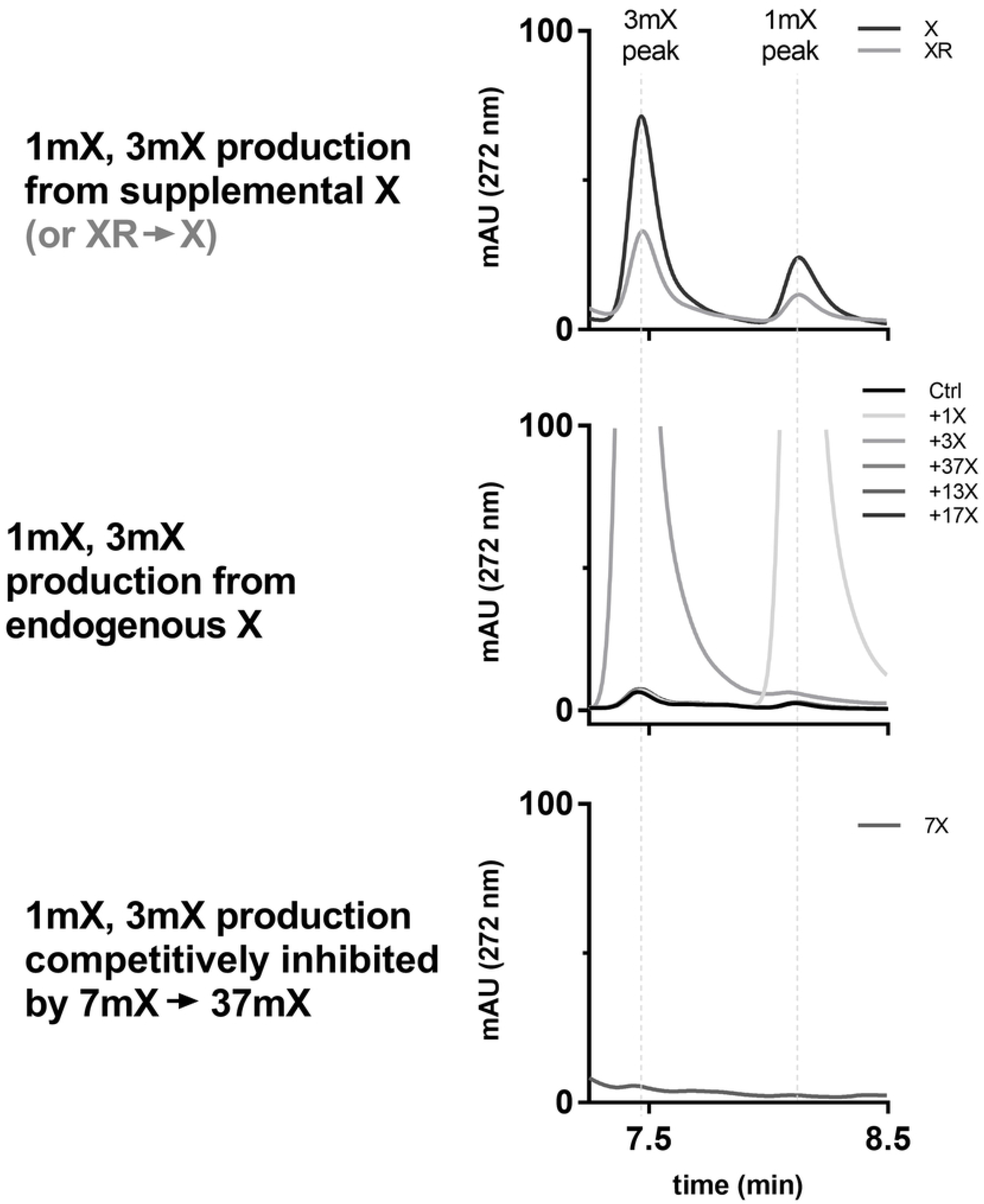
Detail of CsAncCS chromatograms showing 3mX and 1mX peaks for each of added xanthine derivative substrates. Top: with added X (or X produced from XR in situ), 1mX and 3mX are produced. Middle: without an exogenous source of X, 1mX and 3mX are produced in small quantity from endogenous xanthine pools. Bottom: production of 1mX and 3mX from endogenous X is suppressed by competing reaction with 7mX.

Given that we have replicated the high activity observed for predicted ancestral enzymes CsAncCS and PcAncCS2, future efforts towards metabolic engineering of cells for the production of methylxanthines would benefit from a thorough evaluation of the kinetic properties of these under-studied enzymes. This includes enzymes only recently-identified in caffeine producing plants such as PcCS1 and PcCS2 (*Paullinia cupana* caffeine synthase 1 and 2, respectively)[7,13] as well as enzymes from plants which produce methylxanthines (but not necessarily caffeine) such as those from *Theobroma cacao* or *Citrus spp*.[13,31–33] These enzymes as well as their predicted ancestral versions (e.g. predicted TcAncXMT2 from *Theobroma cacao*[32] or CitrusAncXMT2 from *Citrus japonica*[22]) could have desirable properties that are not found in extant enzymes.

### Enzymatic activity of CCS1 can be used as a reporter for juxtacrine cell signaling

Having demonstrated that these enzymes were active in mammalian cells, we wanted to test whether the activity of a methylxanthine synthesis enzyme could be used as a reporter for gene activation. We chose to use an anti-GFP [34] synthetic Notch (synNotch) receptor,[35–39] LaG17-synNotch-TetRVP64, as a means of transducing extracellular binding events to induce production of a reporter enzyme. We designed a reporter construct composed of dTomato and CCS1 linked by a self-cleaving P2A sequence[40] and inserted it into a modified tetracycline-inducible Sleeping Beauty transposon vector to produce the plasmid pSBTet(−ΔrtTA)-GB dTomato-P2A-CCS1. This reporter plasmid would allow induction of both a visible reporter (dTomato) and an enzymatic reporter (CCS1) in the typical Tet-Off fashion [41] that can be triggered by activation of anti-GFP synNotch receptor (**Figure 6**). HEK-293T cells were co-transfected with pSBTet(−ΔrtTA)-GB dTomato-P2A-CCS1, pSBBi-Puro LaG17-synNotch-TetRVP64, and pCMV(CAT) T7-SB100. The transduced cells were selected with puromycin and blasticidin to produce a stable cell line containing both the synNotch receptor and Tet-Off inducible CCS1 reporter circuit. These cells are hereafter called Reporter Cells. K562 suspension cells were lentivirally-transduced to express membrane-bound GFP; these cells are hereafter called Target Cells.

**Figure 6:**
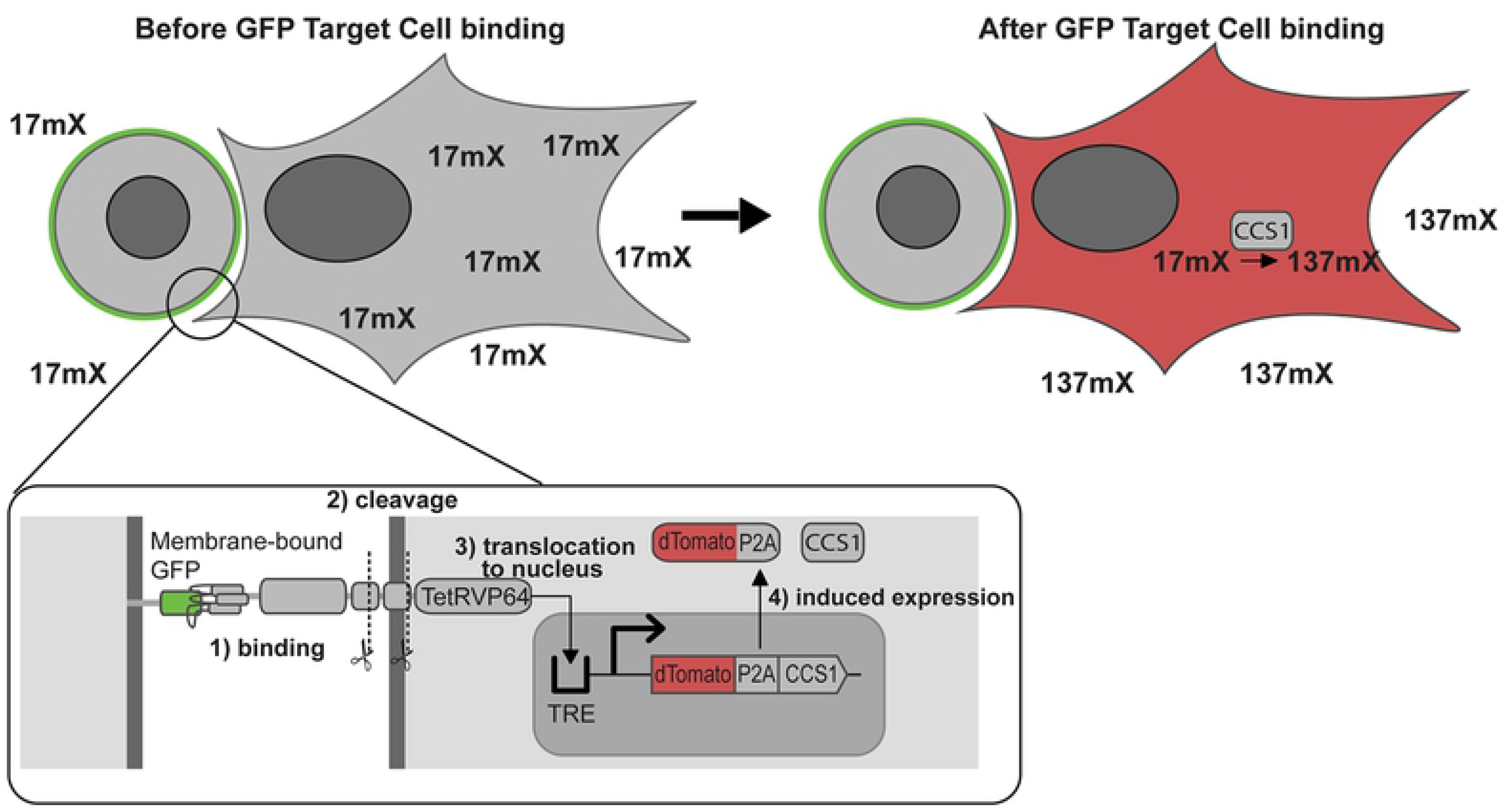
Depiction of how cell-cell contact between Target Cells and Reporter Cells can induce *Coffea* caffeine synthase 1 expression to enable production of caffeine. Inset: the LaG17 anti-GFP binding domain of the synNotch receptor binds surface-expressed GFP in K562 cells.subsequent cleavage events liberate TetRVP64 (tTA). tTa translocates to the nucleus and interacts with the tetracycline response element (TRE) to induce gene expression. The expressed enzyme is capable of converting the supplied 17mX to 137mX (caffeine).

We seeded cell culture flasks with fixed quantities of CCS1 Reporter Cells and variable quantities of Target Cells in culture medium containing 200 μM 17mX. The ratio of Target Cells to CCS1 Reporter Cells ranged from 0:1 to 3:1. After 72 h, the supernatant was harvested and analyzed via HPLC, as before (**Figure 7**). The integrated chromatograms were used in conjunction with a standard curve to determine caffeine concentrations. (**Figure 8**). We found that the measured amount of caffeine increased from 1.0 μM when no GFP Target Cells were present to 14.5 μM when a 3:1 ratio of Target Cells to CCS1 Reporter Cells was used. The low background production of caffeine in the absence of Target Cells is consistent with circuit leakiness seen in other applications of synNotch.[42] We observed a linear dose response relationship between the amount of caffeine produced and the ratio of GFP Target Cells to CCS1 Reporter Cells, although this relationship would be expected to plateau as the ratio of Target Cells to Reporter Cells increases further and synNotch receptors become saturated. While this is admittedly a proof-of-principle experiment, it suggests that the activity of methylxanthine synthesis enzymes could potentially be a viable reporter for induced gene activity in synthetic biology or biochemistry applications.

**Figure 7:**
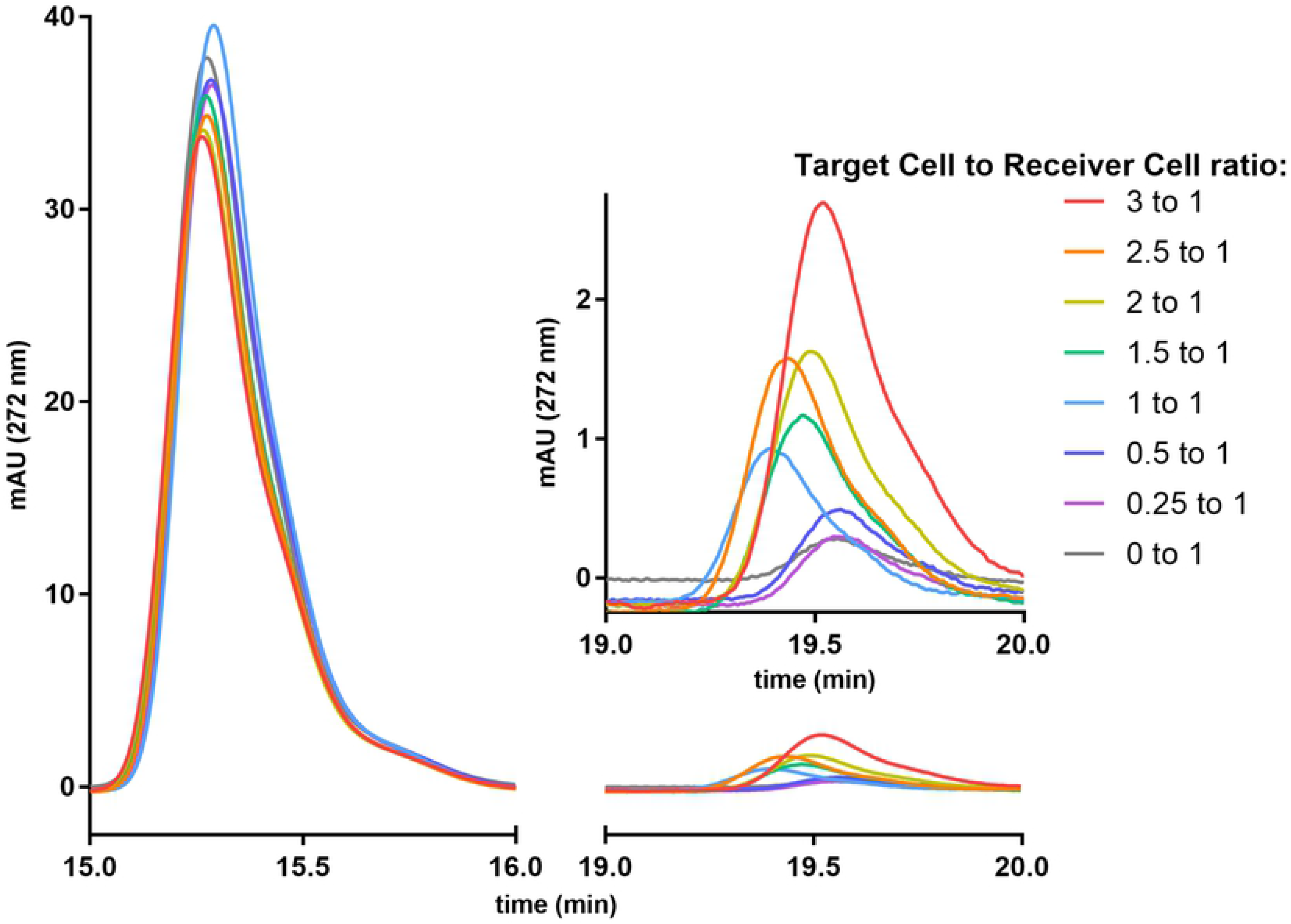
Portions of the chromatograms from coculture of Target Cells with Reporter Cells showing 17mX and 137mX. Left: 17mx peak. Right: 137mX peak. Inset shows more detail of the 137mX peak, highlighting the relationship between the quantity of caffeine produced and the Target to Reporter Cell ratio.

**Figure 8:**
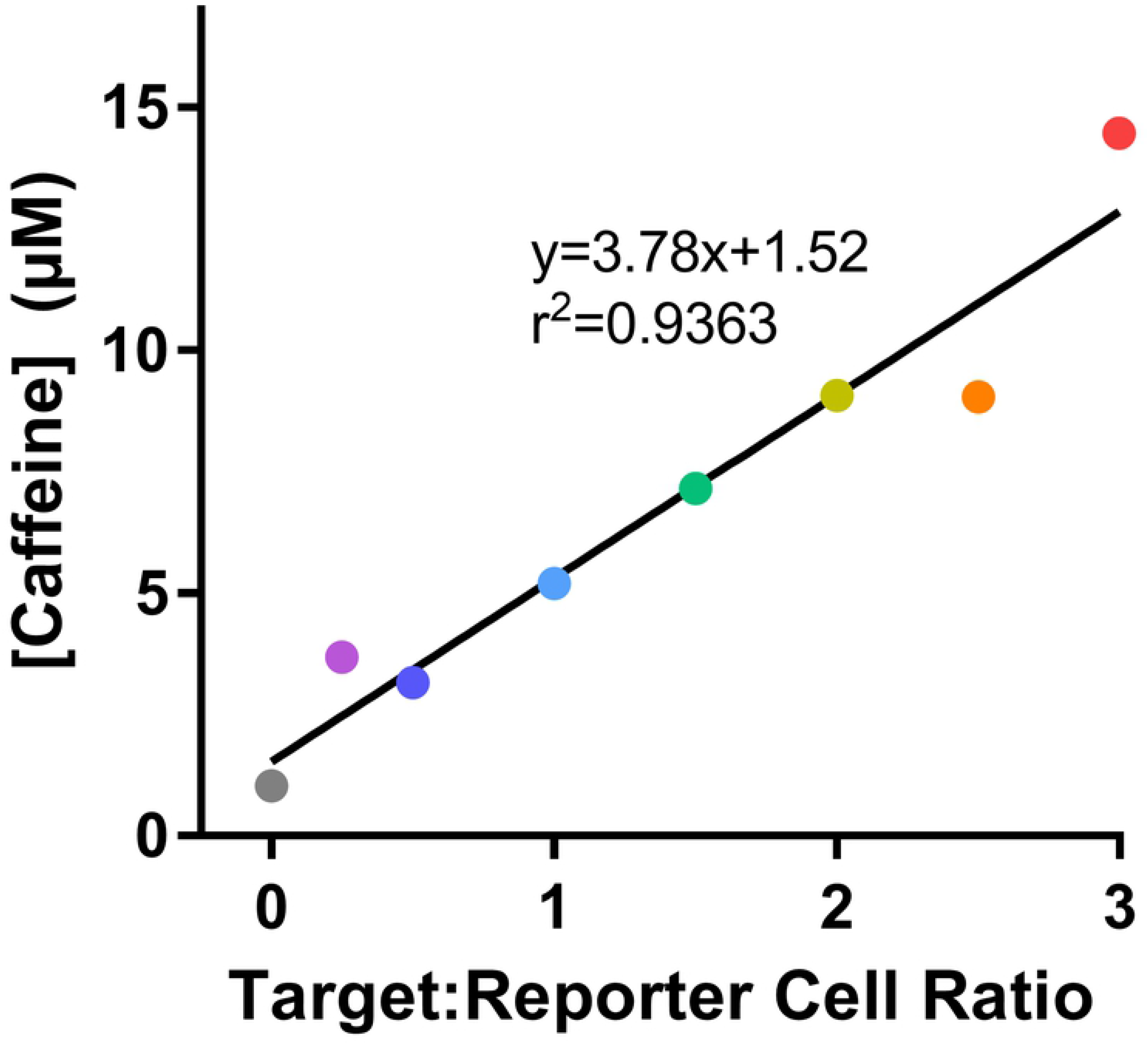
Final caffeine concentration for various Target Cell to Reporter Cell ratios. The linear regression demonstrates dose-response relationship between resultant caffeine concentration and Target Cell to Reporter Cell ratio

### Potential applications for methylxanthine synthesis enzyme expression in mammalian cells

Methylxanthine synthesis enzymes may have potential application for use as a reporter for *in vivo* studies. We suggest that a methylxanthine synthesis enzyme reporter could be useful for longitudinal monitoring of gene activity through testing of urine for methylxanthine metabolites. If CCS1 were used as a reporter enzyme and 7mX were supplied in food, 37mX would be produced upon reporter gene expression. The metabolism of 37mX produces metabolites not produced by 7mX such as 3,7-dimethyluric acid, 3-methyluric acid, and 3-methylxanthine.[43–45] Urine can be easily collected[46], and the quantity of these metabolites could thus be used to assess reporter gene activation. This could even be performed over long periods of time as 7mX has been shown to be non-toxic in both rats and mice.[47,48] This approach also has the benefit of potentially being useable in large animal models as well, where the greater tissue attenuation precludes the use of optically-detectable reporters (i.e. fluorescent and bioluminescent proteins) for longitudinal studies.[49,50] The most promising potential use for these enzymes, however, may be related to the body of work involving synthetic receptors sensitive to caffeine or other methylxanthines.

A small body of previous work has focused on the development of anti-caffeine antibodies, particularly an anti-caffeine camelid heavy-chain-only antibody fragment, aCaffVHH. [51–53] Recent work by Bojar, et al. [21], has demonstrated that aCaffVHH can be fused to proteins that transduce a signal upon dimerization to a produce synthetic receptors that are responsive to caffeine. Although these are described as caffeine receptors, they are not exclusively responsive to caffeine and show some sensitivity to dimethylxanthines; theophylline, paraxanthine, and theobromine all show some activation of the STAT3-based caffeine receptor at 1 μM.[21] Indeed, this pattern of reactivity is observed in binding thermodynamics[52] and in cross reactivity with dimethylxanthines during immunoaffinity chromatography.[51] Previous work developing a water quality immunoassay for caffeine using different monoclonal and polyclonal anti-caffeine antibodies indicated that xanthine produced negligible signal (<0.002% and <0.008% for mAb and pAb immunoassays, respectively).[54] Depending on the relative activation of these receptors with nonmethylated, monomethylated, dimethylated, and trimethylated xanthines, it seems plausible that these receptors could be useful for detecting production of a monomethylated xanthine from a nonmethylated xanthine or a dimethylated xanthine from a monomethylated xanthine.

One could easily imagine a scenario wherein a methylxanthine molecule that is produced in one population of cells (via the enzymes previously discussed) could be used as a diffusible signaling molecule to induce a behavioral change in distant cells using one of the engineered caffeine receptors (**Figure 9**). In this context, methylxanthines could be conceived of as a synthetic paracrine hormone that is largely orthogonal to other signaling pathways. Use of methylxanthines in this manner is comparable with methods used for developing multiplexed intercellular communication with small, organic, signaling molecules derived from ordinary cellular metabolites.[20] This approach could have applications in the study of synthetic tissue development or other contexts where programmed intercellular communication is desired.

**Figure 9: Proposed circuit for cell-cell signaling from a caffeine (or other methylxanthine) producing cell to a detector cell.**

Synthetic caffeine receptor DB326 homodimerizes in the presence of caffeine and activates JAK1 (Janus kinase 1) which activates the phosphorylation and dimerization of STAT3 (signal transducer and activator of transcription 3). The STAT3^P^ dimer can then enable the expression of a STAT3-inducible reporter gene

## Materials and methods

### Materials and chemicals

All chemicals and reagents were of reagent grade or better. Unless otherwise stated, all reagents were purchased from Fisher Scientific (Waltham, MA) or Sigma Aldrich (St. Louis, MO). All solvents for liquid chromatography were HPLC grade or higher and were purchased from Sigma Aldrich or Fisher Scientific. Theobromine, theophylline, and 1-methylxanthine were purchased from Tokyo Chemical Industrial Co., Ltd. (Tokyo, Japan). 3-Methylxanthine was purchased from AK Scientific (Union City, CA). Xanthine was purchased from Chem Impex International (Wood Dale, IL). 7-Methylxanthine was purchased from Carbosynth (San Diego, CA). Caffeine was purchased from Alfa Aesar (Tewksbury, MA).

Dulbecco’s Modified Eagle’s Medium (DMEM), Opti-MEM, HBSS, PBS, TrypLE, polybrene, blasticidin S hydrochloride, and puromycin dihydrochloride were purchased from Thermo Fisher Scientific (Waltham, MA). Heat-inactivated fetal bovine serum (FBS) was purchased from Omega Scientific (Tarzana, CA). Primocin was purchased from Invivogen (San Diego, CA). Cell culture dishes and flasks were purchased from VWR (Radnor, PA). Poly-L-lysine was purchased from Sigma Aldrich (St. Louis, MO). Polythethyleneimine, linear MW 25000, Transfection Grade (PEI) was purchased from Polysciences, Inc. (Warrington, PA). Accutase was purchased from Innovative Cell Technologies (San Diego, CA).

Carbenicillin-containing (100 μg/mL) and kanamycin-containing (50 μg/mL) LB agar plates were purchased from Biopioneer (San Diego, CA). UltraPure Agarose, and LB broth powder were purchased from Thermo Fisher Scientific (Waltham, MA). NEB Stable competent *E. coli* cells were obtained from New England Biolabs (Ipswich, MA). Frozen aliquots of these cells were prepared with the “Mix & Go!” Transformation Kit (Zymo Research, Irvine, CA). All restriction enzymes, Q5 polymerase, HiFi Assembly 2X master mix, T4 DNA ligase, and KLD enzyme mix, were purchased from New England Biosciences (Ipswich, MA). Pre-cast SDS-PAGE gels were purchased from Bio-Rad (Hercules, CA).

### Plasmids

SynNotch plasmids pHR_SFFV_LaG17_synNotch_TetRVP64 and pHR_EGFPligand, were a gift from Wendell Lim (Addgene plasmid # 79128 and 79129, respectively). The Sleeping Beauty transposase plasmid pCMV(CAT) T7-SB100 was a gift from Zsuzsanna Izsvak (Addgene plasmid # 34879). The Sleeping Beauty plasmids pSBBi-RP, pSBBi-GB, pSBBi-Pur, and pSBTet-GB were a gift from Eric Kowarz (Addgene plasmid # 60513, 60520, 60523, and 60504, respectively). Plasmids pMD2.G and pCMV-dR8.2 dvpr were a gift from Didier Trono (Addgene plasmid # 12259 and 8455).

The following genes for methylxanthine synthesis enzymes were ordered from Twist Biosciences (San Francisco, CA) or as gene blocks from Integrated DNA Technologies (Coralville, IA): CaDXMT (AB084125), CaMXMT (MW263197), CaXMT1 (MW263198), CCS1 (MW263199), CsAncCS (MW309842), PcAncCS2 (MW309843), PcCS (BK008796.1), TCS1 (AB031280.1), and TCS2 (MW269521). DNA sequences are available in **S2 File.**

The Sleeping Beauty expression vectors were prepared by use of HiFi Assembly, T4 ligation after restriction digest(s), and/or a Q5 Site-Directed Mutagensis Kit with only slight modifications to manufacturer-provided protocols. Detailed protocols are available in **S3 File.**

The tet-inducible reporter circuit plasmids backbone was prepared by modification of pSBTet-GB to remove constitutive expression of rtTA found in the parent plasmid via site-directed mutagenesis. Normally, the constitutive expression of rtTA allows for inducible gene expression from pSBTet-GB upon the addition of tetracycline. Removal of constitutive rtTA expression means that the inducible protein will only be expressed in the presence of the tetracycline trans-activator derived from a different source. The finished reporter plasmids were then made by HiFi assembly of pSBTet(-ΔrtTA)-GB with dTomato-P2A from pSBBi-RP and CCS1. In this work, these plasmids are always co-expressed with pSBBi-P LaG17-synNotch-TetRVP64 which will liberate its tetracycline trans-activator (i.e. TetRVP64) upon activation of the receptor and thus induce expression of the gene of interest.

The following plasmids are available from Addgene: pSBBi-RP PCCS (#139159), pSBBi-RP TCS1(#139161), pSBBi-RP CCS1 (#139162), pSBBi-RP CaDXMT (#139165), pSBBi-RP CsAncCS (#139167), pSBBi-GB CaXMT1 (#139168), pSBBi-RP PcAncCS2 (#139170), pSBBi-GB TCS2 (#139171), pSBBi-RP CaMXMT (#139172), and pSBBi-Pur LaG17-synNotch-TetRVP64 (#158133).

### Mammalian Cell Culture

#### General Information

HEK-293T (#CRL-3216) and K562 (#CCL-243)cells were purchased from American Type Culture Collection (Manassas, VA). All mammalian cell culture was carried out in a biosafety cabinet using appropriate sterile technique. Cells were counted using a Bio-Rad TC20 cell counter. Adherent cells were detached with TrypLE. Cells were split when cultures reached >90% confluence for adherent cells or when the concentration exceeded 1.5 × 10^6^ cells/mL for suspension cells. Antibiotics were not used routinely and were only used as prophylactic treatment for cell sorting. All routine cell culture media consisted of 10% FBS in DMEM. Cells were tested for mycoplasma every 6 months.

#### Lentivirus production

HEK-293T cells were grown to approximately 90% confluence in a 10-cm dish coated with poly-L-lysine to prevent detachment of cells. Plasmids were diluted to 250 μL in OptiMEM so that the final solution contained 8 μg pMD2.G, 18 μg pCMV-d8.2 dvpr, and 12 μg of pHR_EGFPligand. A polyethyleneimine solution (114 μL of 1 μg/μL) was diluted to 250 μL using OptiMEM. The plasmid and PEI solutions were then combined, gently mixed, and left to incubate at room temperature for 30 minutes. The solution was gently pipetted carefully across the whole surface of the plate and the plate was gently rocked side-to-side to mix. Conditioned media was collected at 24-hour intervals for 3 days, concentrated using a 100 kMW Amicon spin filter and frozen at −80 °C if not used immediately.

#### Lentiviral transduction

A desired quantity of lentivirus concentrate was diluted in DMEM +10% FBS containing 8 μg/mL polybrene. K562 cells were centrifuged, washed once with HBSS, and resuspended with the cell culture medium containing lentivirus and polybrene to a final concentration of 2×10^5^ cells/mL. 2 mL of this cell suspension was added to a well in a 6-well plate. The plate was sealed in two zipper-top bags and centrifuged in a pre-warmed swinging bucket centrifuge for 90 min at 33 °C and 1000 RCF. After incubating the plate for 24 hours at standard cell culture conditions, the medium was replaced with fresh cell culture medium and the cells were cultured as usual, splitting when the concentration exceeded 1×10^6^ cells/mL. After approximately 1 week, the cells were sorted to retain GFP-positive cells by flow-assisted cell sorting.

#### Flow-assisted cell sorting (FACS)

Cells to be sorted were detached using Accutase instead of TrypLE. The detachment was halted by addition of an excess of HBSS or PBS. The cells were pelleted by centrifugation. The supernatant was removed and the cells were washed 3 times with sort buffer (0.5% (w/v) BSA, and 25 mM HEPES in pH 7 PBS). The cell suspension was passed through a 40 μm cell strainer to produce a single-cell suspension. The cells were counted and then diluted to a final concentration of 3 to 7 million cells/mL.

Cells were sorted as the experiment needed using a BD Influx cell sorter (BD Biosciences, San Jose, CA) by staff at the UCSD Human Embryonic Stem Cell Core Facility at Sanford Consortium for Regenerative Medicine (La Jolla, CA). After sorting, they were re-plated at the desired concentration in 50% conditioned medium / 50% fresh medium with Primocin at a final concentration of 100 μg/mL. The Primocin is maintained in culture medium of sorted cells for at least 1 week after sorting to ensure the cells do not become contaminated.

#### Sleeping Beauty transduction and selection

HEK-293T cells were plated in poly-L-lysine-coated 24-well plates and grown until ~90% confluent in 500 μL of medium. Typically, 475 ng of transposon plasmid and 25 ng of pCMV(CAT) T7-SB100 were diluted to 50 μL with Opti-MEM medium. 1.5 μL of Lipofectamine 2000 was added to 50 μL of Opti-MEM. The Lipofectamine 2000 and DNA solutions were mixed gently with a pipette tip and incubated at room temperature for 30 minutes. The prepared DNA lipoplexes were carefully pipetted over the surface of each well and the plate was gently rocked side to side to mix. The cells were left to grow for 24 hours in the incubator.

The next day, cells were inspected on the microscope to check for presence of the fluorescent reporter from the Sleeping Beauty transposon plasmid. If strong fluorescence was not observed, the cells are incubated for an additional 24 hours before re-inspecting. If strong fluorescence was not observed after 48 hours, the cells were discarded, and the transduction was repeated. Once strong fluorescence was observed, the culture was split to a 12-well plate well in 1 mL of medium containing puromycin or blasticidin at the concentrations shown in **Table 3**. Approximately every 3 days, the selection antibiotic concentrations were increased to concentrations shown in the same table. Once the final concentration was reached, the selection antibiotics were maintained for a further 14 days. During this time, the cells populations were expanded and aliquots were frozen on day 14 upon cessation of antibiotic use.

**Table 3:** Antibiotic Selection Concentrations

| Day | [Puromycin]<br>($\mu\text{g/mL}$) | [Blasticidin S]<br>($\mu\text{g/mL}$) |
| --- | --- | --- |
| 1 | 0.5 | 7.5 |
| 4 | 1.0 | 15.0 |
| 7 | 1.5 | 22.5 |
| 10 | 2.0 | 30.0 |

To transduce cells with two plasmids, the same process is carried out except the initial transduction includes two plasmids instead of one. In this case, the total quantity of DNA and lipofectamine were increased to 750 ng and 2.25 μL, respectively. The initial selection was started at the lowest listed concentration of each antibiotic, but the concentration was only increased if it was visually evident that there were enough cells to continue. The antibiotic concentrations were increased, as before, when three days had elapsed or the cell population looked sufficiently large to tolerate the increased selection pressure.

#### Methylxanthine synthesis enzyme substrate activity assay

Cells that constitutively express the various methylxanthine synthesis enzymes were all screened in the same fashion. Cells (0.5 mL of 2×10^5^ cells/mL) in DMEM with 10% FBS were added to each well of a poly-l-lysine-coated 12-well plate. After 24 hours, 1 mL of 300 μM of one of the enzyme substrates was added to each well (final volume = 1.5 mL). If any wells are unused, they were filled 1.5 mL of medium to limit variability in evaporation from plate to plate (**Table 4**). These were incubated for 72 hours in a cell culture incubator. At 72h, the cell medium supernatant was removed in its entirety from each well and transferred to a pre-weighed sample tube. The tube was weighed, and its weight was recorded. The tubes are centrifuged for 2 minutes at 16,000 x RCF and 30 μL used for HPLC analysis. Metabolite peaks on the chromatogram from the diode array detector at 272 nm are positively identified by comparison with known standards. The peaks are integrated using the Agilent Chemstation software and the concentration is determined using a calibration curve of known concentrations for each substance.

**Table 4:** Example plating diagram for enzyme substrate activity assay in a 12-Well Plate

|  |  |  |  |
| --- | --- | --- | --- |
| 200 μM X | 200 μM XR | 200 μM 1mX | Medium |
| 200 μM 3mX | 200 μM 7mX | 200 μM 37mX | Medium |
| 200 μM 13mX | 200 μM 17mX | CTRL | Medium |

The concentration is then corrected to what the concentration would have been without evaporation (i.e. it is multiplied by ratio of the masses of unevaporated medium to recovered medium) so that the semi-quantitative concentration can be more easily compared to the starting concentration. The mass of unevaporated medium used for this was the average of three samples prepared of at the time of plating cells for each experiment. This is necessary because there are differences in evaporative loss across the wells of a 12-well plate and between plates at different positions in a stack.

Experiments with co-culture of TCS2- or CaXMT1-expressing cells were performed similar the standard assay with only minor modifications. TCS2- or CaXMT1-expressing cells (0.25 mL of 2×10^5^ cells/mL) in DMEM with 10% FBS were added to each well. The same amount (0.25 mL of 2×10^5^ cells/mL) of CCS1 cells were added to the co-culture wells and 0.25 mL of DMEM +10% FBS was added to the other wells. After 24 h, 1 mL of 600 μM XR was added to each well. The protocol was otherwise unchanged.

#### Juxtacrine signalling-induced expression of CCS1

For all experiments, HEK-293T cells were doubly transduced (via Sleeping Beauty, as per usual) with pSBBi-P LaG17-synNotch-TetRVP64 and pSBTet(-ΔrtTA)-GB dTomato-P2A-CCS1 and selected with blasticidin and puromycin to produce the CCS1 Reporter Cells. The Reporter Cells were plated at an areal density of 40,000 cells/cm^2^ with a culture medium volume to surface area ratio of 1 mL/5 cm^2^, e.g. a T-25 flask would have 1 million reporter cells seeded and 5 mL of culture medium added. 1,7-Dimethylxanthine was present at a final concentration of 200 μM. The K562 Target Cells, produced by lentiviral transduction and described previously, were seeded simultaneously the desired ratio. This plating scheme can be scaled for 24-well plates, 12-well plates, and T-25 flasks easily. The cell culture medium is harvested at 72 hours and is analyzed by the typical method with HPLC.

### High performance liquid chromatography

Reverse phase high performance liquid chromatography (HPLC) and liquid chromatography with mass spectrometry (LCMS) were performed on an Agilent Infinity 1260 series (Santa Clara, CA) with diode-array detector (DAD) and a single quad mass spectrometer. An Agilent Polaris 5 C18-A column (180Å, 4.6 × 250 mm, 5 μm) or Polaris 5 C18-A (180Å, 10.0 × 250 mm, 5 μm) were used for all separations involving xanthine derivatives. The standard gradient conditions used can be found in the **Table 5**, but this method can be modified as needed to account for differences in column quality over time. If additional elution time was needed, the solvent mix was simply maintained at 83%/17% water/acetonitrile for a desired period after 14 min. The subsequent steps then have their times shifted later by a corresponding amount.

**Table 5:** HPLC method for separation of xanthines

| Time (min) | % Solvent A (H<sub>2</sub>O + 0.1% TFA) | % Solvent B (ACN + 0.1% TFA) |
| --- | --- | --- |
| 0 | 92 | 8 |
| 14 | 83 | 17 |
| 16 | 5 | 95 |
| 20 | 5 | 95 |
| 27.5 | 92 | 8 |
| 36 | 92 | 8 |

Peaks are identified by comparison to known standards prepared using conditioned cell culture medium. Mass spectrometry could sometimes be used to identify peaks, but the high ionic content of the samples likely impaired efficient ionization of the xanthines and produced low intensities. All quantifications were performed by integrating chromatogram peaks from absorbance at 272 nm and calculating the concentration of the substance using a linear regression obtained from a standard curve of known concentrations. Given the culture medium sample composition has a high ionic content, future revisions of the method would likely be improved by use of buffer in solvents A and B instead of pure solvents or involve pre-extraction/desalting of the samples.

## Supporting information

S1 File. HPLC Chromatograms

S2 File. Enzyme Sequences

S3 File. Cloning Protocols

## References

1. Barnes PJ. Theophylline. Am J Respir Crit Care Med. 2013;188: 901–906. doi:10.1164/rccm.201302-0388PP

2. Zoumas BL, Kreiser WR, Martin R. Theobromine and Caffeine Content of Chocolate Products. J Food Sci. 1980;45: 314–316. doi:10.1111/j.1365-2621.1980.tb02603.x

3. Fulgoni VL, Keast DR, Lieberman HR. Trends in intake and sources of caffeine in the diets of US adults: 2001-2010. Am J Clin Nutr. 2015;101: 1081–1087. doi:10.3945/ajcn.113.080077

4. Barone JJ, Roberts H. Human Consumption of Caffeine. Caffeine. Springer Berlin Heidelberg; 1984. pp. 59–73. doi:10.1007/978-3-642-69823-1_4

5. Mitchell DC, Knight CA, Hockenberry J, Teplansky R, Hartman TJ. Beverage caffeine intakes in the U.S. Food Chem Toxicol. 2014;63: 136–142. doi:10.1016/j.fct.2013.10.042

6. Kato M, Mizuno K, Crozier A, Fujimura T, Ashihara H. Caffeine synthase gene from tea leaves. Nature. 2000;406: 956–957. doi:10.1038/35023072

7. Schimpl FC, Kiyota E, Mayer JLS, Gonçalves JFDC, Da Silva JF, Mazzafera P. Molecular and biochemical characterization of caffeine synthase and purine alkaloid concentration in guarana fruit. Phytochemistry. 2014;105: 25–36. Available: https://www.sciencedirect.com/science/article/pii/S0031942214001897

8. Uefuji H, Ogita S, Yamaguchi Y, Koizumi N, Sano H. Molecular Cloning and Functional Characterization of Three Distinct N -Methyltransferases Involved in the Caffeine Biosynthetic Pathway in Coffee Plants. Plant Physiol. 2003;132: 372–380. doi:10.1104/pp.102.019679

9. Mizuno K, Kato M, Irino F, Yoneyama N, Fujimura T, Ashihara H. The first committed step reaction of caffeine biosynthesis: 7-methylxanthosine synthase is closely homologous to caffeine synthases in coffee (Coffea arabica L.). FEBS Lett. 2003;547: 56–60. doi:10.1016/S0014-5793(03)00670-7

10. Kato M, Mizuno K, Fujimura T, Iwama M, Irie M, Crozier A, et al. Purification and characterization of caffeine synthase from tea leaves. Plant Physiol. 1999;120: 579–586. doi:10.1104/pp.120.2.579

11. Mizuno K, Okuda A, Kato M, Yoneyama N, Tanaka H, Ashihara H, et al. Isolation of a new dual-functional caffeine synthase gene encoding an enzyme for the conversion of 7-methylxanthine to caffeine from coffee (Coffea arabica L.). 2003;534: 75–81. doi:10.1016/S0014-5793(02)03781-X

12. Yoneyama N, Morimoto H, Ye CX, Ashihara H, Mizuno K, Kato M. Substrate specificity of N-methyltransferase involved in purine alkaloids synthesis is dependent upon one amino acid residue of the enzyme. Mol Genet Genomics. 2006;275: 125–135. doi:10.1007/s00438-005-0070-z

13. Huang R, O’Donnell AJ, Barboline JJ, Barkman TJ, O’Donnell AJ, Barboline JJ, et al. Convergent evolution of caffeine in plants by co-option of exapted ancestral enzymes. Proc Natl Acad Sci. 2016;113: 10613–10618. doi:10.1073/pnas.1602575113

14. Figueirêdo L, Faria-Campos A, Astolfi-Filho S, Azevedo J. Identification and isolation of full-length cDNA sequences by sequencing and analysis of expressed sequence tags from guarana (Paullinia cupana). Genet Mol Res. 2011;10: 1188–1199. doi:10.4238/vol10-2gmr1124

15. Medrano JF, Cantu D, Hulse-Kemp A, Van Deynze A. The UC Davis Coffea arabica Genome Project. In: Coffea arabica UCDv 0.5. [Internet]. Available: http://phytozome.jgi.doe.gov

16. Denoeud F, Carretero-Paulet L, Dereeper A, Droc G, Guyot R, Pietrella M, et al. The coffee genome provides insight into the convergent evolution of caffeine biosynthesis. Science (80-). 2014;345: 1181–1184. doi:10.1126/science.1255274

17. Xia EH, Zhang H Bin, Sheng J, Li K, Zhang QJ, Kim C, et al. The Tea Tree Genome Provides Insights into Tea Flavor and Independent Evolution of Caffeine Biosynthesis. Mol Plant. 2017;10: 866–877. doi:10.1016/j.molp.2017.04.002

18. Xia E, Li F, Tong W, Yang H, Wang S, Zhao J, et al. The tea plant reference genome and improved gene annotation using long-read and paired-end sequencing data. Sci data. 2019;6: 122. doi:10.1038/s41597-019-0127-1

19. Wei C, Yang H, Wang S, Zhao J, Liu C, Gao L, et al. Draft genome sequence of Camellia sinensis var. sinensis provides insights into the evolution of the tea genome and tea quality. Proc Natl Acad Sci U S A. 2018;115: E4151–E4158. doi:10.1073/pnas.1719622115

20. Du P, Zhao H, Zhang H, Wang R, Huang J, Tian Y, et al. De novo design of an intercellular signaling toolbox for multi-channel cell–cell communication and biological computation. Nat Commun. 2020;11: 1–11. doi:10.1038/s41467-020-17993-w

21. Bojar D, Scheller L, Hamri GC-E El, Xie M, Fussenegger M, Charpin-El Hamri G, et al. Caffeine-inducible gene switches controlling experimental diabetes. Nat Commun. 2018;9: 2318. doi:10.1038/s41467-018-04744-1

22. Huang R. Evolution of Caffeine Biosynthetic Enzymes and Pathways in Flowering Plants. Diss West Michigan Univ. 2017;3169. Available: https://scholarworks.wmich.edu/dissertations/3169

23. Ogawa M, Herai Y, Koizumi N, Kusano T, Sano H. 7-Methylxanthine methyltransferase of coffee plants. Gene isolation and enzymatic properties. J Biol Chem. 2001;276: 8213–8218. doi:10.1074/jbc.M009480200

24. Kowarz E, Löscher D, Marschalek R. Optimized Sleeping Beauty transposons rapidly generate stable transgenic cell lines. Biotechnol J. 2015;10: 647–653. doi:10.1002/biot.201400821

25. Ivics Z, Hackett PB, Plasterk RH, Izsvák Z. Molecular reconstruction of sleeping beauty, a Tc1-like transposon from fish, and its transposition in human cells. Cell. 1997;91: 501–510. doi:10.1016/S0092-8674(00)80436-5

26. Izsvák Z, Ivics Z, Plasterk RH. Sleeping beauty, a wide host-range transposon vector for genetic trensformation in vertebrates. J Mol Biol. 2000;302: 93–102. doi:10.1006/jmbi.2000.4047

27. Jin Z, Maiti S, Huls H, Singh H, Olivares S, Mátés L, et al. The hyperactive Sleeping Beauty transposase SB100X improves the genetic modification of T cells to express a chimeric antigen receptor. Gene Ther. 2011;18: 849–856. doi:10.1038/gt.2011.40

28. Alexiou M, Leese HJ. Purine utilisation, de novo synthesis and degradation in mouse preimplantation embryos. Development. 1992.

29. Li M, Sun Y, Pan SA, Deng WW, Yu O, Zhang Z. Engineering a novel biosynthetic pathway in: Escherichia coli for the production of caffeine. RSC Adv. 2017;7: 56382–56389. doi:10.1039/c7ra10986e

30. Jin L, Bhuiya MW, Li M, Liu XQ, Han J, Deng WW, et al. Metabolic Engineering of Saccharomyces cerevisiae for Caffeine and Theobromine Production. Riezman H, editor. PLoS One. 2014;9: e105368. doi:10.1371/journal.pone.0105368

31. Jones PG, Allaway D, Gilmour DM, Harris C, Rankin D, Retzel ER, et al. Gene discovery and microarray analysis of cacao (Theobroma cacao L.) varieties. Planta. 2002;216: 255–264. doi:10.1007/s00425-002-0882-6

32. O’Donnell AJ. Evolutionary Convergence of the Caffeine Biosynthetic Pathway in Chocolate Followed Duplication of a Constrained Ancestral Enzyme. Master’s Theses, West Michigan Univ. 2015;602. Available: https://scholarworks.wmich.edu/masters_theses

33. Koyama Y, Tomoda Y, Kato M, Ashihara H. Metabolism of purine bases, nucleosides and alkaloids in theobromine-forming Theobroma cacao leaves. Plant Physiol Biochem. 2003;41: 977–984. doi:10.1016/j.plaphy.2003.07.002

34. Fridy PC, Li Y, Keegan S, Thompson MK, Nudelman I, Scheid JF, et al. A robust pipeline for rapid production of versatile nanobody repertoires. Nat Methods. 2014;11: 1253–1260. doi:10.1038/nmeth.3170

35. Roybal KT, Rupp LJ, Morsut L, Walker WJ, Mcnally KA, Park JS, et al. Precision Tumor Recognition by T Cells With Combinatorial Antigen-Sensing Circuits. Cell. 2016;164: 770–779. doi:10.1016/j.cell.2016.01.011

36. Toda S, Blauch LR, Tang SKY, Morsut L, Lim WA. Programming self-organizing multicellular structures with synthetic cell-cell signaling. Science (80-). 2018;361: 156–162. doi:10.1126/science.aat0271

37. Gordley RM, Bugaj LJ, Lim WA. Modular engineering of cellular signaling proteins and networks Introduction: why design and engineer signaling proteins? Curr Opin Struct Biol. 2016;39: 106–114. doi:10.1016/j.sbi.2016.06.012

38. Roybal KT, Lim WA. Synthetic Immunology: Hacking Immune Cells to Expand Their Therapeutic Capabilities. Annu Rev Immunol. 2017;35: 229–53. doi:10.1146/annurev-immunol

39. He L, Huang J, Perrimon N, Potter CJ, Venken K. Development of an optimized synthetic Notch receptor as an in vivo cell–cell contact sensor. Proc Natl Acad Sci. 2017; 201703205. doi:10.1073/pnas.1703205114

40. Szymczak-Workman AL, Vignali KM, Vignali DAA. Design and construction of 2A peptide-linked multicistronic vectors. Cold Spring Harb Protoc. 2012;7: 199–204. doi:10.1101/pdb.ip067876

41. Gossen M, Bujard H. Tight control of gene expression in mammalian cells by tetracycline-responsive promoters. Proc Natl Acad Sci U S A. 1992;89: 5547–5551. doi:10.1073/pnas.89.12.5547

42. Roybal KT, Williams JZ, Morsut L, Rupp LJ, Kolinko I, Choe JH, et al. Engineering T Cells with Customized Therapeutic Response Programs Using Synthetic Notch Receptors. Cell. 2016;167: 419–432.e16. doi:10.1016/j.cell.2016.09.011

43. Miller GE, Radulovic LL, Dewit RH, Brabec MJ, Tarka SM, Cornish HH. Comparative theobromine metabolism in five mammalian species. Drug Metab Dispos. 1984;12: 154–160.

44. Arnaud MJ. Pharmacokinetics and metabolism of natural methylxanthines in animal and man. Handbook of Experimental Pharmacology. 2011. doi:10.1007/978-3-642-13443-2_3

45. Cornish HH, Christman AA. A study of the metabolism of theobromine, theophylline, and caffeine in man. J Biol Chem. 1957;228: 315–323. Available: http://www.ncbi.nlm.nih.gov/pubmed/13475320

46. Kurien BT, Everds NE, Scofield RH. Experimental animal urine collection: A review. Lab Anim. 2004;38: 333–361. doi:10.1258/0023677041958945

47. Singh H, Sahajpal NS, Singh H, Vanita V, Roy P, Paul S, et al. Pre-clinical and cellular toxicity evaluation of 7-methylxanthine: an investigational drug for the treatment of myopia. Drug Chem Toxicol. 2019;0: 1–10. doi:10.1080/01480545.2019.1635615

48. Singh H, Singh H, Sahajpal NS, Paul S, Kaur I, Jain SK. Sub-chronic and chronic toxicity evaluation of 7-methylxanthine: a new molecule for the treatment of myopia. Drug Chem Toxicol. 2020;0: 1–12. doi:10.1080/01480545.2020.1833904

49. Troy T, Jekic-McMullen D, Sambucetti L, Rice B. Quantitative comparison of the sensitivity of detection of fluorescent and bioluminescent reporters in animal models. Mol Imaging. 2004;3: 9–23. doi:10.1162/153535004773861688

50. Darne C, Lu Y, Sevick-Muraca EM. Small animal fluorescence and bioluminescence tomography: A review of approaches, algorithms and technology update. Phys Med Biol. 2014;59. doi:10.1088/0031-9155/59/1/R1

51. Franco EJ, Sonneson GJ, DeLegge TJ, Hofstetter H, Horn JR, Hofstetter O. Production and characterization of a genetically engineered anti-caffeine camelid antibody and its use in immunoaffinity chromatography. J Chromatogr B Anal Technol Biomed Life Sci. 2010;878: 177–186. doi:10.1016/j.jchromb.2009.06.017

52. Sonneson GJ, Horn JR. Hapten-Induced Dimerization of a Single-Domain VHH Camelid Antibody. Biochemistry. 2009;48: 6693–6695. doi:10.1021/bi900862r

53. Ladenson RC, Crimmins DL, Landt Y, Ladenson JH. Isolation and Characterization of a Thermally Stable Recombinant Anti-Caffeine Heavy-Chain Antibody Fragment. Anal Chem. 2006;78: 4501–4508. doi:10.1021/ac058044j

54. Carvalho JJ, Weller MG, Panne U, Schneider RJ. A highly sensitive caffeine immunoassay based on a monoclonal antibody. Analytical and Bioanalytical Chemistry. 2010. pp. 2617–2628. doi:10.1007/s00216-010-3506-1

